# Accumulation of defense systems in phage resistant strains of *Pseudomonas aeruginosa*

**DOI:** 10.1101/2022.08.12.503731

**Authors:** Ana Rita Costa, Daan F. van den Berg, Jelger Q. Esser, Aswin Muralidharan, Halewijn van den Bossche, Boris Estrada Bonilla, Baltus A. van der Steen, Anna C. Haagsma, Ad C. Fluit, Franklin L. Nobrega, Pieter-Jan Haas, Stan J.J. Brouns

## Abstract

Prokaryotes encode multiple distinct anti-phage defense systems in their genomes. However, the impact of carrying a multitude of defense systems on phage resistance remains unclear, especially in a clinical context. Using a collection of antibiotic-resistant clinical strains of *Pseudomonas aeruginosa* and a broad panel of phages, we demonstrate that defense systems contribute substantially to defining phage host range and that overall phage resistance scales with the number of defense systems in the bacterial genome. We show that many individual defense systems are specific to phage genera, and that defense systems with complementary phage specificities co-occur in *P. aeruginosa* genomes likely to provide benefits in phage-diverse environments. Overall, we show that phage-resistant phenotypes of *P. aeruginosa* with at least 19 phage defense systems exist in the populations of clinical, antibiotic-resistant *P. aeruginosa* strains.

## Introduction

Bacteriophage predation imposes a strong evolutionary pressure on bacteria to evolve mechanisms to defend against phage infection (*1*). These defense mechanisms include modification of cell surface receptors (*2, 3*) and intracellular defenses (*4–6*) such as CRISPR-Cas (*7, 8*) and Restriction-Modification (RM) (*9, 10*).

More recently, dozens of previously unknown anti-phage immune systems have been discovered. In most instances, they were identified based on the observation that immune systems often cluster in defense islands (*11–16*). The presence and composition of these defense islands vary among individual strains (*4, 15, 17*), and strongly contribute to phage-host co-evolution in natural populations (*1*). The presence of multiple variable defense systems in bacterial genomes raises the important question what the impact is of all these immune systems on overall phage resistance of bacterial pathogens.

To address this question, we assembled a set of 32 clinical, antibiotic-resistant *Pseudomonas aeruginosa* strains and compiled a custom panel of 28 phages from 12 phylogenetic groups. We then analyzed phage infectivity and adsorption of the strains across the panel. This revealed that intracellular phage defense mechanisms are an important determinant of the phage susceptibility of *P. aeruginosa*, and that strains rich in phage defense systems are inherently more resistant to phage infection. Five strains contained a large number (13–19) of anti-phage defense systems and displayed an extended phage-resistant phenotype, in addition to having an extended drug-resistant (XDR) phenotype. Our data further revealed that defense systems can be specific to certain phage families, and that the activity of these individual defense systems in model strains can often predict the resistance of clinical strains to the same phages. Additionally, we have found that some combinations of defense systems with complementary phage specificity often co-occur in *P. aeruginosa* genomes and may provide phage defense with broader phage specificity. Overall, our findings have implications for our understanding of phage defense and potentially for the development of phage-based antibacterial therapeutics, as antibiotic-resistant strains with an extended phage-resistant phenotypes are present in clinical settings.

## Results

### Defense systems are abundant and diverse in clinical *P. aeruginosa* strains

We selected a set of 32 antibiotic-resistant clinical strains covering the diversity of defense systems in the *P. aeruginosa* species as a whole. The *P. aeruginosa* genomes from the RefSeq database carry 71% (119/167) of the known defense system subtypes (**Fig. 1a, Table S1**), which is in line with recent observations that *P. aeruginosa* has a diverse arsenal of anti-phage defense (*17*). The defense systems found in the RefSeq genomes resembled the defense arsenal in our clinical isolates (**Fig. 1b, Table S2**), and the number of defense systems per genome ranged between 1 and 19 systems for both datasets (**Fig. S1c**,**d**). Additionally, we assessed the phylogenetic distribution of our collection using a maximum likelihood analysis (*18*), which revealed the distribution of the clinical isolates across the two main phylogroups, with 23 strains in the largest phylogeny group 1 and 9 strains in phylogeny group 2 (**Fig. S1**). We found that the RefSeq genomes belonging to phylogeny group 2 carried slightly higher number of defense systems compared to those of phylogeny group 1 (median phylogenetic group 1 = 7, median phylogenetic group 2 = 9, p = 0.0004), but these differences are not observed in the strains present in our collection (p = 0.612) (**Fig. S1**). Overall, we demonstrate that *P. aeruginosa* genomes encode multiple different anti-phage defense systems, and establish that our collection of clinical isolates is phylogenetically diverse and covers the range of defense systems both in types and numbers per strain as observed in *P. aeruginosa* as a whole.

**Fig. 1.**
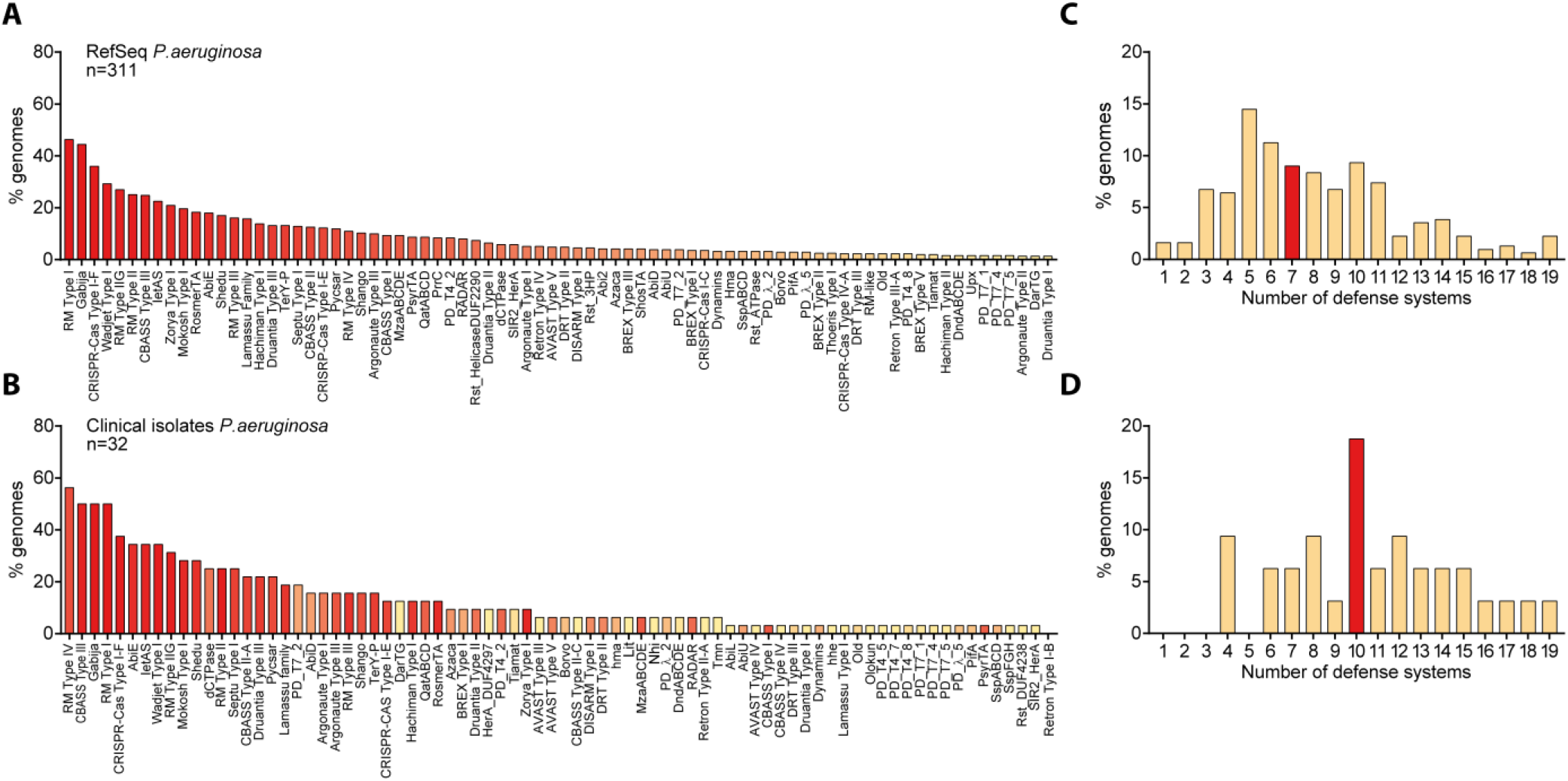
Defense systems are abundant and diverse in *P. aeruginosa* strains. **(A)** Diversity of defense systems found in the genomes of 311 *P. aeruginosa* strains from the RefSeq database, organized and colored in a gradient from most (left) to least (right) abundant. Only the most prevalent defense systems are shown (see **Table S1** for the full list of defense systems). **(B)** Diversity of defense systems found in the genomes of 32 clinical isolates of *P. aeruginosa* from our collection. Systems are organized from most (left) to least (right) prevalent and colored according to the abundance in (A). All defense systems found in the clinical strains are shown. **(C)** Number of defense systems per genome in *P. aeruginosa* strains from the RefSeq database. **(D)** Number of defense systems per genome in *P. aeruginosa* strains from our collection of clinical isolates. For (C) and (D) the median number of defense systems is shown in red.

### Phage resistance correlates with the number of defense systems

To obtain a relevant panel of phages for these clinical isolates, we used a subset of 22 *P. aeruginosa* strains as hosts to enrich and isolate different phages from sewage water. We obtained a total of 27 phages (**Table S3**), consisting of *Caudoviricetes* (dsDNA tailed phages), including 13 podophages (5 *Autographiviridae*, 3 *Bruynoghevirus*, 1 *Schitoviridae*, 1 *Zobellviridae*, and 3 unassigned phages), 4 siphophages (1 *Samunavirus*, 1 *Casadabanvirus*, 1 *Mesyanzhinovviridae*, and 1 *Detrevirus*), and 10 myophages (9 *Pbunavirus* and 1 *Phikzvirus*, a Jumbo myophage related to nucleus-forming *Pseudomonas* phage phiKZ (*19*)). To broaden the diversity of our phage panel beyond dsDNA phages, we additionally included *Fiersviridae* PP7 (ssRNA phage). We then used vConTACT2 (*20*) to assess the taxonomic diversity of our phage panel, and found that it represents 9 out of the 16 phage clusters observed in *P. aeruginosa* phages overall, thus indicating a diverse representation of phages (**Fig. S2**). To determine the effect of the defense systems on the susceptibility of the clinical strains to our panel of phages, we first assessed the ability of the phages to infect the strains, or to only adsorb to their cell surface without infecting. Out of a total of 924 phage-host combinations (28 phages times 33 hosts, including PAO1), 630 phage-host combinations did not result in infection (**Fig. 2a**). We hypothesize the non-infected phenotype can occur in two ways: either the phage fails to adsorb to the cell surface (i.e., no receptor), or phage propagation is unsuccessful, possibly due to anti-phage defense. Based on adsorption assay data, we found that a large proportion out of the 630 non-infection cases showed adsorption. More precisely, 68% (429) of the phage-host combinations showing no infection exhibited adsorption when considering an adsorption threshold of 50% (**Fig. 2a**), and 32% (201) when adopting a more conservative threshold of 90% (**Fig. S3a**). These results indicate internal defense mechanisms could play a role in preventing infection in a substantial fraction of strains where no infection was observed.

**Fig. 2.**
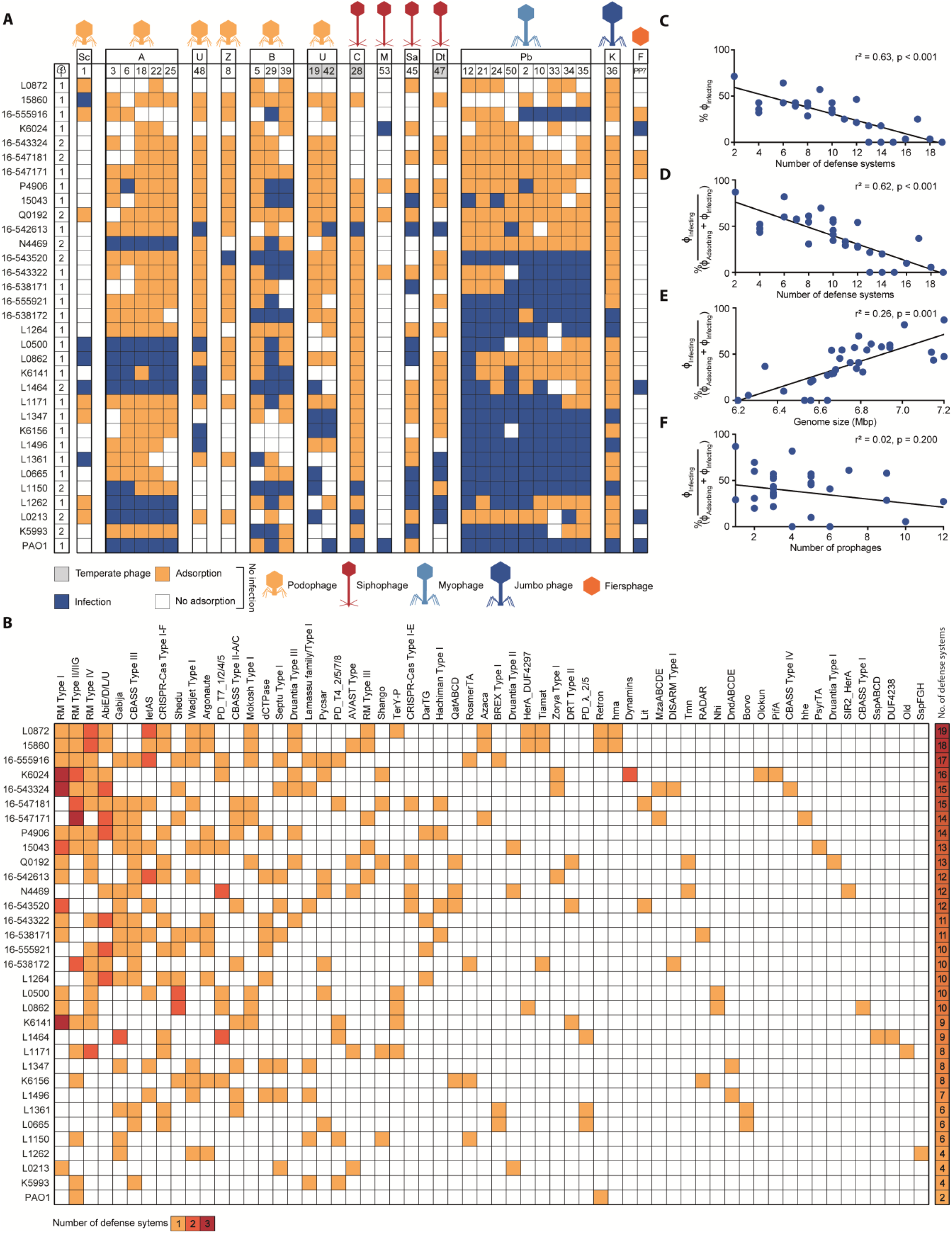
Innate and adaptive defense systems correlate with phage resistance. **(A)** Host range of phages against 32 clinical isolates of *P. aeruginosa* and strain PAO1. Phages are clustered by phylogeny (**Table S3**). Phage-bacteria interactions are depicted as infection (blue), adsorption (>50%) but no infection (orange), or no interaction (white). Letters above the phage numbers indicate family or genus (for phages unassigned to a family): A, *Autographiviridae*; B, *Bruynoghevirus*; C, *Casadabanvirus*; Dt, *Detrevirus*; F, Fiersviridae; K, *Phikzvirus*; M, *Mesyanzhinovviridae*; Pb, *Pbunavirus*; Sa, *Samunavirus*; Sc, *Schitoviridae*; Z, *Zobellviridae*; U, unassigned. **(B)** Defense systems found in the *P. aeruginosa* clinical isolates. The number of instances of each defense system type per strain is indicated in yellow, orange, or red for 1, 2, or 3 respectively. The total number of defense systems found per strain is indicated in a heatmap bar on the right. A complete list of the defense systems found in the clinical isolates can be found in **Table S2**. **(C)** Linear regression analysis of how number of defense systems found in the *P. aeruginosa* clinical isolates and PAO1 correlate with the levels of phage resistance. **(D)** Linear regression analysis of how number of defense systems correlates with the levels of phage resistance calculated as the percentage of adsorbing phages that can establish a productive infection (% ϕ_Infecting_ / (ϕ_Adsorbing_+ϕ_Infecting_)). **(E)** Linear regression analysis of how genome size correlates with phages that can establish a productive infection. **(F)** Linear regression analysis of how number of prophages correlates with phages that can establish a productive infection. In panels (B), (C), (D), and (E), r^2^ represents R-squared, a goodness-of-fit measure for the linear regression models.

We observed that some of the *P. aeruginosa* strains (L0872, Q0192, 16-543324, and 16-547171), which encoded more defense systems than average, exhibited complete resistance to our phage panel (**Fig. 2a,b**) and questioned which factor is the main driver of phage resistance among the number of defense systems, genome size, and number of prophages. To assess this, we performed a multiple linear regression analysis including these factors, which showed that defense systems are the only relevant indicator of phage resistance (r^2^ = 0.63 and p < 0.001 for correlation with % infecting phages; r^2^ = 0.62 and p < 0.001 for correlation with % adsorbing phages that can establish a productive infection (% ϕ_Infecting_ / (ϕ_Adsorbing_+ϕ_Infecting_)) (**Fig. 2c-f, Fig. S3b-e**). The correlation between defense systems and phage resistance also remained robust and significant when considering a conservative adsorption threshold of 90% (r^2^ = 0.35 and p < 0.001) (**Fig. S3f-h**), or when only one representative phage per family or genus was included (# defense systems r^2^ = 0.48 and p < 0.001, genome size r^2^ = 0.11 and p = 0.033, # prophage r^2^ = -0.02 and p = 0.6256) (**Fig. S3i-l**). Although the number of prophages did not show significant correlation with phage resistance, we did observe mechanisms of superinfection exclusion(*21*) for strains L1361 and L1496, which contain one prophage each that likely provides protection against closely related temperate phages ϕPa47 (100% pident, 100% coverage) and ϕPa42 (100% pident, 89% coverage), respectively.

Overall, our findings suggest that the number of defense systems in *P. aeruginosa* is associated with phage resistance. This observation is best exemplified by the five *P. aeruginosa* strains (L0872, 16-543324, 16-547171, 16-547181, Q0192) with 13-19 defense systems (**Fig. 2a,b**) that were found to be resistant to the complete phage panel and to our attempts of phage isolation using wastewater from different sources. Strikingly, these strains are also resistant to multiple antibiotics (**Table S2**), indicating that clinical isolates can exhibit simultaneous resistance to a broad range of antibiotics and phages.

In summary, we show that phage resistant strains of *P. aeruginosa* have accumulated phage defense systems in their genome, suggesting that phage defense systems could be a contributing factor of the phage sensitivity of the host.

### Adaptive immunity targets temperate phages

Half (16/32) of the clinical strains contain adaptive immune systems in the form of CRISPR-Cas Type I-F (12 strains) and Type I-E (4 strains). To investigate the contribution of CRISPR-Cas to phage resistance we identified all spacers targeting our phage panel and assessed their potential effect on the phage host range. We detected 70 spacers (43 unique) across 16 strains matching our phage panel (**Fig. S4a**), among which 54 are interference-proficient (i.e., spacers with matching protospacer adjacent motif (PAM) and protospacer) and 16 are priming-proficient (i.e., spacers with a ±1 slipped PAM(*22*) or up to 5 protospacer mutations (*23, 24*)) (**Table S4**). The majority of the spacers (65) originate from CRISPR-Cas Type I-F systems, with only five spacers from CRISPR-Cas Type I-E systems (strains 15-547181, N4469, and 15043, all targeting ϕPa28). Interestingly, 66 of the 70 spacers target temperate phages (ϕPa19, ϕPa28, ϕPa42, and ϕPa47), and only four spacers match a virulent phage (ϕPa8). This is in line with previous findings that spacers of *P. aeruginosa* mostly match temperate phages (*25*). Our data on phage infection and adsorption reveal that in 83% (25/30) of cases with matching I-E (4/4) and I-F (21/26) spacers, the targeted phage was unable to infect (**Fig. S4a**). Out of the five cases where the protective effects of matching spacers were not observed (**Fig. S4a**), one was linked to the presence of an Acr (ϕPa42 infecting 16-542613) (**Fig. S4b, Table S5**).

Overall, our results suggest that CRISPR-Cas Type I-E and I-F may contribute to resistance of the clinical strains against temperate phages but plays a minor role against the vast majority of virulent phages in the panel because they are not targeted. Although the catalog of spacers may evolve through CRISPR adaptation, the current set of spacers alone does not explain the observed infection profiles.

### Innate defense systems provide anti-phage activity against specific phage families

To understand the contribution of individual innate defense systems to broad-spectrum phage immunity, we inserted 14 individual defense systems (**Fig. S5a**) from *P. aeruginosa* clinical isolates into the low-copy plasmid pUCP20(*26*), under their native promoters. The plasmids were introduced into *P. aeruginosa* strain PAO1, which is infected by 18 phages of our panel. We validated that the defense systems represent no burden or toxicity to cell growth (**Fig. S5b**), and subsequently assessed the defense-containing PAO1 strains for changes in phage susceptibility using efficiency of plating (EOP), infection dynamics, and bacterial culture collapse assays.

The EOP (**Fig. 3a**) and infection dynamics (**Fig. S6**) assays show that 9 out of 14 defense systems exhibit activity against the phage panel, with most of them displaying specificity towards phage families. For example, both TerY-P and Zorya are active against podophages in our panel, but TerY-P seems to be restricted to *Autographiviridae* while Zorya also targets podo *Bruynoghevirus*, sipho *Casadabanvirus*, and *Fiersviridae*. QatABCD and RADAR are both strong defenses against myo *Pbunavirus* and target members of different families of siphophages (*Casadabanvirus* and *Mesyanzhinovviridae*, respectively). This specificity is consistent with previous reports of defense systems sensing conserved phage proteins to trigger an immune response (*27–30*). CBASS Type III-C displays the broadest protection in our set, acting against podo *Autographiviridae*, sipho *Mesyanzhinovviridae* and *Casadabanvirus*, myo *Pbunavirus*, and Jumbo *PhiKZvirus*. This suggests that CBASS Type III-C employs a more general mechanism that relies on conserved phage features shared among different families, or that its effector is activated by other cellular responses, such as a general stress response. Interestingly, Zorya Type I is the only defense system among those tested that prevents infection of ssRNA phage PP7 (*Fiersviridae*), suggesting also that the effector may be activated by general cellular responses to phage infection. Additionally, CBASS Type III-C and AVAST Type V systems provide robust (>10^5^-fold) protection against infection by phiKZ-like, nucleus-forming Jumbo phage ϕPa36 (**Fig. 3a,b**). Both systems have been reported to act via altruistic cell death upon sensing of a specific phage protein (*27, 31, 32*), a strategy that (like RNA-targeting CRISPR-Cas systems (*33*)) circumvents the nuclear shell defense used by Jumbo phages to overcome DNA-targeting defense systems (*34–36*). The bacterial culture collapse assays provided additional information about the protective effect of the defense systems in liquid culture. TerY-P and Zorya Type I protect the cell population at both low and high phage multiplicity of infection (MOI) (**Fig. 3b, Fig. S7**), and most cells survive infection monitored using microscopy with propidium iodide as an indicator of membrane permeability and cell death (**Fig. 3c**). For QatABCD, RADAR, Druantia Type III, AVAST Type V, CBASS Type III-C, II-A, and II-C, a culture collapse is observed when the phage is introduced at high MOI. In summary, our findings indicate that individual defense systems display phage-targeting activity that is mostly specific to particular phage families. While the majority of the defense systems could provide protection against phages at low concentration, only two were efficient against phage at high densities.

**Fig. 3.**
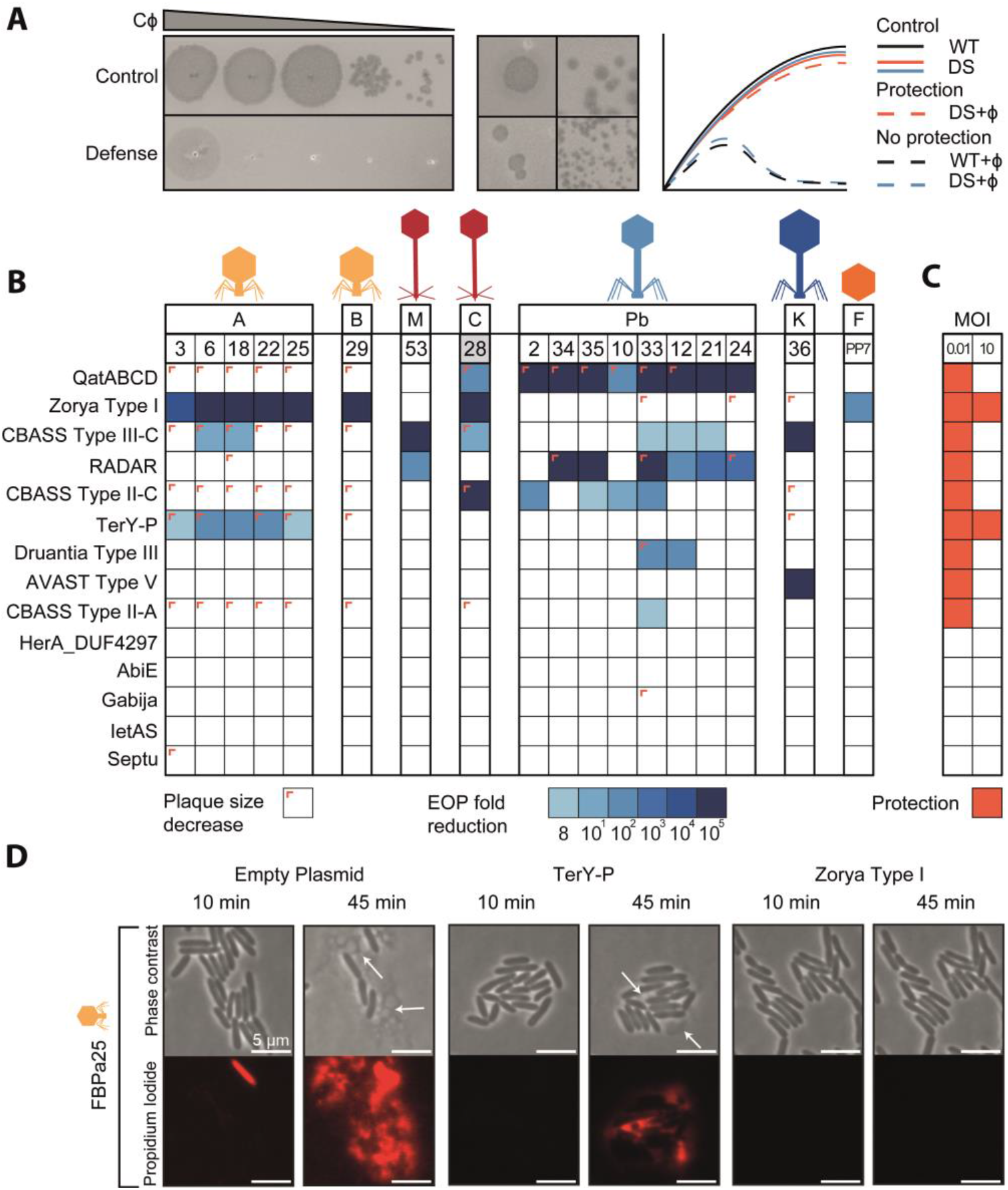
Defense systems provide genera-specific anti-phage activity. **(A)** Representation of the assays used to assess the effect of individual defense systems on phage infectivity. On the left, an example of fold decrease in ϕPa33 phage infectivity caused by the defense system QatABCD. In the middle, examples of phage plaque size decrease observed for *Autographiviridae* ϕPa3 and *Casadabanvirus* ϕPa28 with QatABCD cells. On the right, illustrations of growth curves obtained from liquid culture collapse assays for control strains (WT) and strains containing individual defense systems (DS) with and without phage infection, showing cases of protection (orange) and no protection (blue). **(B)** Efficiency of plating (EOP) of phages in PAO1 containing individual defense systems. EOP was determined as the fold decrease of phage titer in the strain with the defense system compared to the titer obtained in the strain without the defense system. Plaque size reductions are indicated as a colored corner. **(C)** Liquid culture collapse assays of PAO1 containing individual defense systems infected with phages, as compared to a control (WT) without a defense system. Results are shown as a summary of the effects observed when infecting the cultures with a multiplicity of infection (MOI) of 10 or 0.01. A protective effect is represented as dark orange. **(D)** Time-lapse phase contrast and fluorescence images of PAO1 cells containing individual defense systems infected with phage ϕPa25. Cells were stained with propidium iodide to visualize permeabilization of the cell membrane due to cell death. TerY-P and Zorya Type I cells survive phage infection, although some cell death is still observed for TerY-P, consistent with the EOP results in (B).

### Linking native infection profiles with protection patterns of individual systems

To understand the overall phage protection observed in genetically inaccessible clinical strains (**Fig. 2**) in relation to the effect of individual phage defense systems (**Fig. 3**), we initially assessed the phage infectivity levels in these strains using efficiency of plating assays (**Fig. S8a**). We then compared the phage susceptibility profiles of the clinical strains with those of PAO1 strains equipped with a single defense system. Our findings indicate that in 84% of the cases, the phage susceptibility profile of the clinical strains aligns with the expected profile (**Fig. S8b)**. This is especially evident for Zorya Type I, TerY-P, Druantia Type III, AVAST Type V, and CBASS Type II-A. The most notable disparity in the results was observed for RADAR and CBASS Type II-C, as they were providing protection against *Pbunavirus* in PAO1 and not in the clinical strains (ϕPa34 and ϕPa35 for RADAR; ϕPa2, ϕPa10, ϕPa33, and ϕPa35 for CBASS Type II-C).

To assess if known phage-encoded anti-defenses impact the phage infectivity profile of the clinical strains, we searched phage genomes for anti-defense genes including anti-RM (*37–39*), anti-CBASS (*28, 40*), anti-Pycsar (*40*), anti-TIR-STING (*41*), and anti-AVAST (*27*) proteins (**Fig. S4b, Table S5**). We focus here specifically on the anti-defenses against the defense systems introduced in PAO1, which include anti-CBASS and anti-AVAST. Our search identified one phage-encoded anti-defense gene, an anti-CBASS Type II (*acbII*) in phage ϕPa48. The impact of *acbII* on the phage host range is not clear since ϕPa48 can only infect two (K6141 and L1496) out of the nine strains that carry CBASS Type II (**Fig. 2a,b**). This outcome is possibly due to other defense systems that target ϕPa48 in these strains.

In addition, we found an *acbII* in strains L1347 and 16-547171, which carry the CBASS Type II-C and II-A systems, respectively. This gene appears to facilitate phage infection in four out of six cases (**Fig. 2a, Fig. S4b**). The suppression of *acbII* expression or the presence of other defense systems may be the reasons why in two cases the bacteria can resist phages despite having an anti-defense gene.

Overall, our findings underscore the intricate nature of phage susceptibility in natural settings, which is likely influenced by the interaction between different defense and anti-defense mechanisms present in both the strain and the phage.

### Co-occurrence of defense systems with complementary specificities

Based on the observation that some defense systems provide distinct genera-specific anti-phage activities (**Fig. 3**), we hypothesized that combinations of defense systems may be advantageous for cells by providing a wider protective range, and would be a conserved feature in bacterial genomes to efficiently achieve broader antiviral specificity. To test this hypothesis, we assessed the co-occurrence (i.e. presence in the same genome) of defense systems in *P. aeruginosa* genomes in the RefSeq database (n = 311), while taking phylogeny into account (*42*).

We found multiple defense system co-occurrences (147 out of 1317 (11%) combinations tested, Bonferroni-corrected binomial exact test statistic with a p-value < 0.01), seven of which involved defense systems with anti-phage activity in this study (**Table S6**). Of these, six combinations have complementary phage specificity, including: i) Druantia Type III (*Pbunavirus*) and TerY-P (*Autographiviridae*), ii) Druantia Type III and Zorya Type I (*Autographiviridae*, *Bruynoghevirus*, *Casadabanvirus*, *Fiersviridae*), iii) Druantia Type III and AVAST Type V (*Phikzvirus*), iv) AVAST Type V (*Phikzvirus*) and QatABCD (*Casadabanvirus*, *Pbunavirus*), v) AVAST Type V and Zorya Type I (*Autographiviridae*, *Bruynoghevirus*, *Casadabanvirus*, *F*), and vi) TerY-P (*Autographiviridae*) with CBASS Type II (*Casadabanvirus*, *Pbunavirus*). Druantia Type III and CBASS Type II have overlapping specificity for *Pbunavirus*, with CBASS Type II adding specificity to *Casadabanvirus*.

Overall, our analysis suggests that defense systems with complementary anti-phage activity co-occur at a probability higher than by chance in *P. aeruginosa* genomes and could provide an advantage for bacterial survival in phage diverse environments.

## Discussion

Bacterial strains carry numerous distinct phage defense systems in their genomes (*17*). We found that *P. aeruginosa* strains carry at least 71% of all currently known defense systems, making this species a versatile bacterial model to study phage immunology. Using a diverse set of clinical isolates of *P. aeruginosa*, we observe that strains that have accumulated defense systems in their genome display broad and robust immunity against phage infection.

By testing the activity of 14 defense systems against our phage panel, we observed that the majority (7 out of 9 active systems) of the defense systems tested in PAO1 are capable of protecting the cell population at low phage concentration (MOI <1), but not at high concentration (MOI ≥ 1). This phenotype could be caused by the defense system being overwhelmed at high phage concentration, or by death or dormancy of the infected cell (*43*) which serves as a means of protecting the cell population through kin selection. The latter phenomenon has been reported for CBASS (*44*) and for RADAR (*45, 46*), and is now proposed for QatABCD, Druantia Type III, and AVAST Type V. In addition, we show that defense systems have anti-phage specificity that is often linked to the phage family or genus, suggesting that defense systems operate using conserved properties within phage phylogenetic groups, as shown previously for certain defense systems (*27, 30*). However, CBASS Type III-C and Zorya Type I in our study were effective against multiple distinct phage families, suggesting that these defense systems employs a more general sensing mechanism or that their effector is activated by other cellular responses. While Zorya remains largely uncharacterized, the current knowledge of CBASS activation suggests a variety of phage sensing strategies, including the recognition of peptides (HORMA domain) and dsDNA binding (cyclase) in *Escherichia coli* CBASS Type III-C (*32*), binding of structured phage RNA in *Staphylococcus schleiferi* CBASS Type I-B (*47*), and phage-driven depletion of folate-like molecules in *Vibrio cholerae* CBASS Type II (*48*).

Interestingly, we found that several pairs of the tested defense systems, which exhibit complementary anti-phage specificities, co-occur in *P. aeruginosa* strains. We expect that these combinations of defense systems could provide a broader range of phage protection through the complementary phage specificities of each individual defense system. The complementary activities of naturally co-occurring defense systems have also been found to enable resuscitation from defense system-induced bacterial dormancy (R-M and CRISPR-Cas) (*49*) and to prevent plasmid dissemination in *Vibrio cholerae* El Tor strains (DdmABC and DdmDE) (*50*), the latter proposing that defense system cooperation might play a role in bacterial pathogenicity.

The strong correlation found between number of defense systems and phage resistance further indicates that multiple defense system combinations are beneficial to cover the whole range of predating phages. The importance of the number of defense systems in determining phage resistance is further evidenced by the high levels of phage-resistance of five of our clinical isolates that encode between 13 to 19 defense systems. Attempts to isolate phages from diverse wastewater samples against defense-rich strains or to transform plasmid DNA proved more difficult, again pointing to the strains’ inherent ability to defend well from incoming threats.

Importantly, our findings demonstrate the significance of the individual defense systems in predicting susceptibility of *P. aeruginosa* strains to phages. However, factors such as genetic context, interactions among defense systems (*49, 51–53*), the presence of unknown defense systems and anti-defense mechanisms, will affect the final outcome of phage infection. Further research is required to enhance the predictive accuracy of genomic analysis, which could prove beneficial for ecological and evolutionary studies.

Altogether, our results show that while phage host range has traditionally been linked to receptor-associated factors (*54*), the number of defense systems is also a strong indicator of the susceptibility of cells to phage in *P. aeruginosa*. Naturally occurring *P. aeruginosa* clinical strains with a large number of defense systems show an increased resistance to phages and may be selected for upon more widespread use of therapeutic phages. Therefore, monitoring the evolution and spread of phage-resistant clinical pathogens, and selecting or engineering phages with anti-defense properties may become instrumental to combat antimicrobial resistance using phage.

## Materials and Methods

### Bacteria

A set of 22 clinical isolates of *P. aeruginosa* provided by the University Medical Center Utrecht (UMCU) was used for phage isolation and 32 for characterization of host range (**Table S2**). The antibiotic susceptibility of the strains was established using the broth microdilution method outlined by EUCAST for determining the minimal inhibitory concentration and interpreted according to the EUCAST 2023 breakpoints (www.eucast.org). *Escherichia coli* strain Dh5α was used to clone plasmid pUCP20 with individual defense systems. *P. aeruginosa* strains containing pUCP20 with individual defense systems were constructed from *P. aeruginosa* strain PAO1. All bacterial strains were grown at 37 °C in Lysogeny Broth (LB) with 180 rpm shaking for liquid cultures, or in LB agar (LBA) plates for solid cultures. Strains containing plasmid pUCP20 were grown in media supplemented with 100 µg/ml of ampicillin (for *E. coli*) or 200 µg/ml of carbenicillin (for *P. aeruginosa*).

### Bacteriophages

Phages used in this study are described in **Table S3**. All phages were isolated from sewage water. Approximately 1 ml of sewage sample was added to 20 ml of LB, inoculated with 100 µl of overnight cultures of each *P. aeruginosa* clinical isolate, and incubated overnight at 37 °C with 180 rpm shaking. Samples were centrifuged at 3,000 × *g* for 15 min and filter-sterilized (0.2 µm PES). The phage-containing supernatant was serially diluted in LB and spotted onto double-layer agar (DLA) plates of the isolation strains for the detection of phages. Single plaques with distinct morphologies were picked with sterile toothpicks and spread with sterile paper strips onto fresh bacterial lawns. The procedure was repeated until a consistent plaque morphology was obtained. Phages from purified plaques were then produced in liquid media with their respective host, centrifuged, filter-sterilized, and stored as phage lysates at 4 °C. For EOP and liquid infection assays (see below), phage stocks were obtained from lysates prepared on PAO1 and their concentration normalized to ≈1x10^8^ pfu/ml. Additional efforts were made to isolate phages for the *P. aeruginosa* clinical isolates that exhibited the highest phage-resistance. This involved using sewage water from various sources and following the enrichment procedure outlined above, but with individual strains instead of mixtures.

### Phage host range

Phages were 10-fold serially diluted in LB and spotted onto DLA plates containing each of the 32 *P. aeruginosa* clinical strain used for phage characterization (**Table S2**). The plates were incubated overnight at 37 °C and the phage plaques were observed to distinguish productive infection (lysis with individual phage plaques formed) from lysis from without (*55*) (lysis without individual phage plaques). Efficiency of plating of phages in each clinical strain was determined by comparing phage titer to that obtained in PAO1 (for phages that infect this strain) or in the clinical strain with the highest phage titer (for phages that cannot infect PAO1).

### Adsorption assays

Early-exponential (optical density at 600 nm, OD_600_ ≈ 0.3) cultures of the *P. aeruginosa* clinical isolates were added in triplicates to the wells of 96-well plates. Phages were added to these cultures at an MOI of 0.01 and incubated at 37 °C with 100 rpm shaking for 15 min. The plates were centrifuged and a sample of the supernatant was taken, 10-fold serially diluted, and plated onto DLA plates of PAO1 to determine the titer of phages that did not adsorb to the clinical strain. A control plate in which phages were added to LB was used to determine the total phage concentration. The concentration of adsorbed phages was determined by subtracting non-adsorbed phage concentration from the total phage concentration in the suspension. The percentage of adsorbed phages was calculated as the ratio between adsorbed phages and total phages. Phages were considered to adsorb when over half of their population on average adhered to the cells.

### Extraction of phage DNA and bacterial DNA

Phage DNA was extracted using phenol-chloroform. For this, 5 mL of each phage lysate at >10^9^ pfu/mL were treated with 1 µg/mL of DNase I and RNase for 30 min. Ethylenediaminetetraacetic acid (EDTA), proteinase K, and sodium dodecyl sulfate (SDS) were added to the sample at final concentrations of 20 mM, 50 µg/mL, and 0.5% respectively, and the samples were incubated at 56 °C for 1 h. The samples were then mixed with an equal volume of chloroform and centrifuged at 3,000 × *g* for 10 min. The aqueous phase was recovered and the procedure was repeated sequentially with a 1:1 mixture of phenol:chloroform, and chloroform. The resulting aqueous phase was mixed with 0.1 volume of sodium acetate 3M (pH 5) and 2.5 volumes of ice-cold absolute ethanol and incubated at -20 °C overnight. The extracted DNA was pelleted at 14,000 × *g* for 15 min and washed in ice-cold 70% ethanol, before re-suspending in ultrapure water. Bacterial genomic DNA was extracted using the GeneJET Genomic DNA Purification kit (Thermo Fisher). The quality and quantity of extracted phage and bacterial DNA were estimated using a NanoPhotometer and a Qubit fluorometer, respectively.

### Phage genome sequencing

For samples sequenced at Beijing Genomics Institute (BGI) (**Table S3**), the phage genomic DNA was fragmented by Covaris 55 µL series Ultrasonicator, and used to construct DNA nanoball (DNB)-based libraries by rolling circle replication. DNA was sequenced using the BGI MGISEQ-2000 platform (BGI Shenzhen, China) with a paired-end 100 nt strategy, generating 4.6-19.2 Gb sequencing data for each sample. For phage samples sequenced in-house, phage DNA was fragmented by Covaris M220 Focused-Ultrasonicator, and libraries were prepared using the NEBNext Ultra II DNA Library Prep Kit. Size distribution was checked on an Agilent D1000 Screen Tape System, and the libraries were pooled equally and spiked with approximately 5% of the PhiX control library. The pooled library was sequenced with an Illumina MiSeq using the MiSeq Reagent Nano Kit v2 (500-cycles). For samples sequenced at the Microbial Genome Sequencing Center (MiGS, Pittsburgh, PA, USA), sample libraries were prepared using the Illumina DNA Prep kit and IDT 10 bp UDI indices, and sequenced on an Illumina NextSeq 2000, producing 2x151 bp reads. Demultiplexing, quality control, and adapter trimming were performed with bcl-convert (v3.9.3). Reads obtained for all samples were assembled using Unicycler v0.5.0 (*56*). For samples sequenced in-house, the control PhiX was manually removed from the assembled contigs using Bandage (*57*).

### Bacterial genome sequencing

For samples sequenced at BGI (**Table S2**), the bacterial genome was fragmented by Covaris 55 µL series Ultrasonicator and used to construct paired-end libraries with an insert size of 200-400 bp. Bacterial genomes were sequenced on the BGISEQ-500 (MGI, BGI-Shenzhen) platform, generating 1.4-2.0 Gb sequencing data for each sample with a sequencing depth >100x. Reads were checked for contamination using kraken2 (*58*) and only considered for further analysis if >90% of the reads identified as *P. aeruginosa*. Quality control of the raw data was performed using FastQC (*59*) with default parameters. For samples sequenced at MiGS, sequencing was performed as described above for phages. Reads obtained for all samples were assembled using Unicycler and the assembly quality was assessed using assembly-stats.v1.0.1 (https://github.com/sanger-pathogens/assembly-stats) and BUSCO.v4 (*60*) (pseudomonodales_odb10), and the GC% was calculated using bioawk (https://github.com/lh3/bioawk). The sequencing depth was calculated using minimap2 (*61*) and samtools mpileup (*62, 63*).

### Bacteria genome annotation and phylogenomics

Bacterial genomes of the clinical strains were annotated using Prodigal (*64*). The genomes were used to determine the multi-locus sequence type (MLST) of the strains using the PubMLST website (https://pubmlst.org/) (*65*), and the serotype using the *Pseudomonas aeruginosa* serotyper PAst (https://github.com/Sandramses/PAst). A total of 311 complete *P. aeruginosa* genomes were downloaded from RefSeq on February 2022. A phylogenetic tree was constructed using Parsnp (*66*) with *P. aeruginosa* strain PAO1 (NC_002516.2) as the reference genome. Phylogeny groups were determined as previously described (*18*). The number of complete prophages present in RefSeq and clinical strains was predicted with virsorter2 v2.2.4(*67*), checkv v1.0.1 (*68*) (end_to_end with checkv-db-v1.0) and a second round of virsorter2 v2.2.4, following the protocol described in https://www.protocols.io/view/viral-sequence-identification-sop-with-virsorter2-5qpvoyqebg4o/v3. Superinfection exclusion was considered when the prophage and temperate phage shared a nucleotide similarity of pident >90% and coverage > 85%.

### Phage genome annotation, taxonomy and phylogenomics

Phage genomes were annotated using the RAST server (*69*), the start of the phage genome was determined using PhageTerm (*70*), and partial genes were manually verified and removed. The phage lifestyle was predicted using PhageAI (*71*). Phages from our collection were classified taxonomically using GRAViTy (*72*). Phage diversity was evaluated using vConTACT2 (*20*) with the default settings and the ProkaryoticViralRefSeq94-Merged database, specifically selecting for *P. aeruginosa* phage genomes. The output of vConTACT2 was visualized in a circular layout using Cytoscape (*73*).

### Detection of defense systems in bacterial genomes

Defense systems were detected in the *P. aeruginosa* genomes of RefSeq and clinical isolates with PADLOC-DB v1.4.0 (*74*), DefenseFinder (*17*), and the HMMs with completeness rules and thresholds as applied in Gao *et al* (2020) (*75*). In addition, the representative sequences provided by Rousset *et al* (2023) (*76*) were used to search for the defense system Detocs described in this work. Homology searches were performed via blastp (*77*) (> 0.7 subject length / query length < 1.5 ; 0.7 > query coverage < 1.3; evalue < 1e-9). Systems were considered complete when all genes were present without more than 2 genes in between. In case of discrepancies between the algorithms, we considered the output reporting the most hits. For PADLOC, we excluded defense systems of the “other” categories. For DefenseFinder, we excluded results of defense systems that were not discriminated into subtypes, e.g. BREX.

In addition, a manual search of the neighborhood of the defense systems identified by the algorithms led to the identification of a variant of the TerY-P system that contained all three genes and corresponding functional domains of the original system (*75*). The new TerY-P sequences were used to search for this variant in the bacterial genomes using blastp with evalue < 2.34e-29 and pident > 30. Systems were considered complete when all genes were present with less than 3 genes in between.

Coinfinder (*42*) was used for detecting the co-occurrence of defense systems in the *P. aeruginosa* genomes of the RefSeq database, using the Parsnp (*66*) phylogenetic tree as input.

### Detection of CRISPR-Cas I-F and I-E spacers targeting phages from our collection

Spacers were detected in the bacterial genomes using CRISPRDetect (*78*), and were mapped to our phage collection using blastn (word size = 8; evalue = 1; query coverage > 90; pident > 90; no gaps; maximum of 1 mismatch allowed). The non-target strand PAM (5’-CC for I-F, 5’-AAG for I-E) was manually checked, with a +1 or -1 PAM slippage allowed for I-F (*22*). Spacers with a matching PAM and protospacer were categorized as interference-proficient, while spacers with a PAM slippage or up to five protospacer mutations (with correct PAM) were categorized as priming-proficient spacers.

### Detection of anti-defense genes in bacteriophage and bacterial genomes

Acrs were detected using AcrFinder (*79*). For the detection of anti-RM (*ardA* (*37*), *klcA* (*38*), *ardB*, *ocyA*, *ocr*, *darA*, *darB* (*39*)), anti-CBASS Type I (*acb1*) (*40*), anti-CBASS Type II (*acbII*) (*28*), anti-Pycsar (*apyc*) (*40*), anti-TIR-STING (*41*), and anti-AVAST (*lidtsur-6*, *lidtsur-17*, *forsur-7*, *penshu1-7*, *usur-3*, *smaarsur-6*, and *mellemsur-6*) (*27*) genes, we first searched for *P. aeruginosa* homologs using PSI-BLAST (*80*) (maximum of 3 runs with 500 sequences; coverage > 60%, pident > 20%). Homolog functionality was checked using HMMer (*81*) and HHpred (*82*). *P. aeruginosa* homologs were only found for anti-genes *acb1* and *acbII*. These homologs were searched for in our phage and bacterial genomes with the use of blastp (evalue < 10^-8 ; pident > 30 ; coverage > 60% ; 2.0 < subject length / query length > 0.5). For genes with no *P. aeruginosa* homologs, we created an HMM from the multiple alignment file obtained from the PSI-BLAST search above, using hmmbuild v3.3.2 (*81*) with default settings. These HMMs were used to search for the anti-defense genes in our phage and bacterial collections (evalue < 10^-6). All hits obtained were checked for the presence of the expected functional domains by HMMer and HHpred.

### Cloning of defense systems in PAO1

Defense systems were amplified from *P. aeruginosa* strains using the primers indicated in **Table S7** with Q5 DNA Polymerase (New England Biolabs), in reactions that added regions of homology to plasmid pUCP20. PCR products were run on 1% agarose gels and bands of the desired size were excised and cleaned using the Zymoclean Gel DNA Recovery Kit. Plasmid pUCP20 (pEmpty, **Table S8**) was digested with BamHI and EcoRI, treated with FastAP (Thermo Scientific), and cleaned with the Zymo DNA Clean & Concentrator Kit. Each defense system was cloned into pEmpty using the NEBuilder HiFi DNA Assembly Master Mix, and transformed into chemically competent NEB^®^ 5-alpha Competent *E. coli* following the manufacturer’s instructions. Plasmids were extracted using the GeneJET Plasmid Miniprep kit, confirmed by sequencing (Macrogen, primers in **Table S7**), and electroporated into PAO1 as previously described (*83*). Briefly, an overnight culture of PAO1 was centrifuged at 16,000 × *g* for 2 min at room temperature, and the pellet was washed twice and resuspended in 300 mM of sucrose. The suspension was mixed with 100-500 ng of plasmid DNA and electroporated at 2.5 kV in a 2 mm gap electroporation cuvette. Cells were recovered in LB for 1-2 h at 37 °C and plated in LBA plates supplemented with 200 µg/ml of carbenicillin.

### Efficiency of plating

The 10^8^ pfu/ml phage stocks were 10-fold serially diluted in LB and the dilutions were spotted onto DLA plates of PAO1, or DLA+carbenicillin plates of PAO1 with pEmpty or PAO1 with individual defense systems following the small plaque drop assay (*84*). The phage dilution that resulted in countable phage plaques was used in double-layer overlay plaque assays (*85*) with PAO1, PAO1 with pEmpty, or PAO1 with the defense systems. The anti-phage activity of the systems was determined as the fold reduction in phage plaques in comparison to the number of plaques obtained in the PAO1:pEmpty control. The diameter of the phage plaques was measured to determine differences in plaque size caused by the defense systems.

### Infection dynamics of phage-infected cultures

Bacterial cultures of PAO1 with pEmpty or with individual defense systems at an OD_600_ ≈ 0.1 were infected with phage at an MOI <1. The cultures were incubated at 37 °C with rocking, and samples were taken at 0h, 2h, 4h, and 6h to measure phage concentration. The sample was centrifuged at 3,000 × *g* for 5 min, and the phage-containing supernatant was 10-fold serially diluted and spotted onto DLA plates of PAO1 to estimate phage concentration.

### Liquid culture collapse assays

Overnight grown bacteria were diluted to an OD_600_ of approximately 0.1 in LB media. The cell suspension was distributed into the wells of 96-well plates, and phages were added at MOIs of 10, 0.1, 0.01, and 0.001). Assays were performed in triplicates. The plates were incubated at 37°C in an Epoch2 microplate spectrophotometer (Biotek) for OD_600_ measurements every 10 min for 24h, with double orbital shaking.

### Fluorescence microscopy of defense systems

Exponentially growing (OD_600_ ≈ 0.3) cultures of PAO1 strains containing pEmpty or the defense systems were infected with phage at an MOI of ≥ 3, and the phage was adsorbed for 10 min at 37 °C. Cells were centrifuged at 9,000 × *g* for 1 min, and the cell pellet was re-suspended in 5 µL of 1 µM of propidium iodide. The stained cells were spotted onto 1% agarose pads (*86*), and visualized using a Nikon Eclipse Ti2 inverted fluorescence microscope equipped with a 100× oil immersion objective (Nikon Apo TIRF; 1.49 numerical aperture). Time-lapse phase-contrast (CD Retiga R1) and fluorescence images (after excitation with a 561 nm laser 2000 609/54 bandpass filter, EM-CCD Andor iXON Ultra 897) were acquired every 5 min using Metamorph.

### Statistical analysis

Unless stated otherwise, data are presented as the mean of biological triplicates ± standard deviation. All correlation analysis were determined by linear (multiple) regression models using the lm function of R, and p-values were adjusted with the Bonferroni post-hoc test. The significance of differences between phylogenetic groups was determined using the Kruskal-Wallis test with Dunn’s post-hoc test, while the differences in infection dynamics were determined by two-way ANOVA followed by Sidak’s multiple comparison test. For all statistical analysis, a significance level of 0.05 was used.

## Acknowledgements

We thank Linyi Gao and Feng Zhang (Massachusetts Institute of Technology) for kindly providing the HMMs for the detection of defense systems, and Wenchen Song and Minfeng Xiao (BGI-Shenzhen) for sequencing part of the clinical isolates and phages used in this study. We would like to thank Wim de Leeuw and Han Rauwerda (University of Amsterdam, Swammerdam Institute for Life Sciences, MAD/RB&AB) for the use of the Crunchomics computer cluster. We also thank members of the Brounslab for the many discussions and ideas that improved our work.

## Author contributions

Conceptualization: SJJB

Methodology: SJJB, FLN, ARC, DFB

Software: DFB, AM

Formal Analysis: ARC, DFB, JQE, AM, BAS, HB

Investigation: ARC, DFB, JQE, AM, HB, BEB, ACH

Visualization: ARC, DFB, JQE, HB, AM

Data Curation: ARC, DFB, JQE

Writing – Original Draft: ARC, DFB, JQE

Writing – Review & Editing: SJJB

Resources: PJH, ACF, FLN

Funding Acquisition: SJJB

## Data and materials availability

All data are available in the main text or the supplementary materials, and at https://github.com/BrounsLab-TUDelft/Pseudomonas_Defence_Systems. All original code has been deposited at Github. Raw data and assembled bacterial and phage genomes are available from Genbank, Bioproject PRJNA817167 (individual accession numbers are listed in Supplementary Tables 2 and 3). All unique bacterial strains, phages, and plasmids generated in this study are available from the corresponding author upon request.

## Supplementary material

**Fig. S1.** Phylogenomic analysis of *Pseudomonas aeruginosa* strains.

**Fig. S2.** Diversity of the phage panel used in this study.

**Fig. S3.** Linear regression analysis of correlation between different variables and phage resistance in *Pseudomonas aeruginosa*.

**Fig. S4.** Role of adaptive immunity and anti-defense genes in defining the phage host range in *Pseudomonas aeruginosa*.

**Fig. S5.** Individual defense systems cloned in *Pseudomonas aeruginosa* strain PAO1.

**Fig. S6.** Effect of individual defense systems on phage infection dynamics in *Pseudomonas aeruginosa* PAO1.

**Fig. S7.** Effect of individual defense systems on bacterial growth during phage infection.

**Fig. S8.** Predictive value of defense system activity in PAO1 to phage susceptibility of the clinical strains.

**Table S1.** Matrix of defense systems identified in the 311 RefSeq genomes of *Pseudomonas aeruginosa*.

**Table S2.** Features of the clinical isolates of *Pseudomonas aeruginosa* used in this work, including defense system presence.

**Table S3.** Features of the *Pseudomonas aeruginosa* phages used in this study.

**Table S4.** List of CRISPR-Cas Type I-F and I-E interference-proficient and priming-proficient spacers found in the clinical isolates of *Pseudomonas aeruginosa* to target phages from our collection.

**Table S5.** Anti-defense genes found in our collection of *Pseudomonas aeruginosa* clinical strains and phages.

**Table S6.** Co-occurrence of defense systems in the *Pseudomonas aeruginosa* genomes of the RefSeq database identified by Coinfinder.

**Table S7.** List of primers used in this work.

**Table S8.** List of plasmids used in this work.

