## Supplementary material for "Accumulation of defense systems in phage resistant strains of *Pseudomonas aeruginosa*": Fig. S

**This PDF file includes:**

Figs. S1 to S8

**Other Supplementary Materials for this manuscript include the following:**

Tables S1 to S8

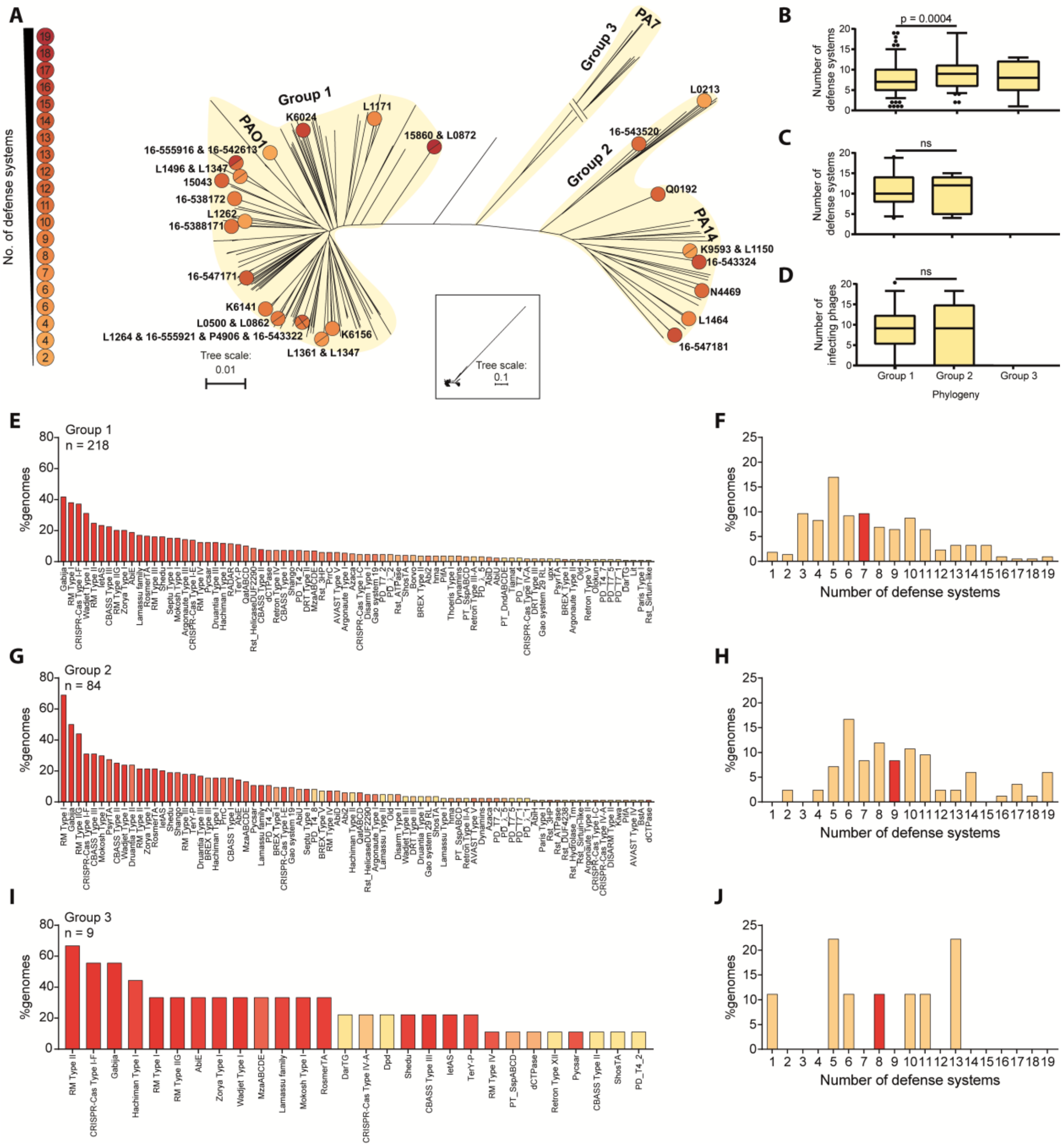


**Fig. S1. Phylogenomic analysis of *Pseudomonas aeruginosa* strains. (A)** Maximum likelihood phylogenetic tree of *P. aeruginosa* strains built using core genome SNPs mapped to the reference strain PAO1 (NC_002516.2) using Parsnp. Circles and squares indicate the positions of clinical strains in phylogenetic groups 1 and 2, respectively, and are colored based on the number of defense systems present in each strain. **(B)** Comparison of the number of defense systems found in the RefSeq *P. aeruginosa* genomes among phylogenetic groups. **(C)** Comparison of defense systems in the genomes of the 32 clinical strains among phylogenetic groups. **(D)** Comparison of the number of infecting phages in the genomes of the clinical strains among phylogenetic groups. For panels (B), (C), and (D), statistical analysis was determined using the Kruskal-Wallis test with Dunn's post-hoc test, with a significance value of 0.05. **(E)** Diversity of defense systems and **(F)** number of defense systems per genome in *P. aeruginosa* strains from the RefSeq database of phylogeny group 1. **(G)** Diversity of defense systems and **(H)** number of defense systems per genome in RefSeq *P. aeruginosa* of phylogeny group 2. **(I)** Diversity of defense systems and **(J)** number of defense systems per genome in RefSeq *P. aeruginosa* of phylogeny group 3. In panels (E), (G), and (I), the defense systems are organized from most (left) to least (right) prevalent and colored according to the abundance in Fig. 1a. In panels (F), (H), and (J), the median number of defense systems is shown in red.

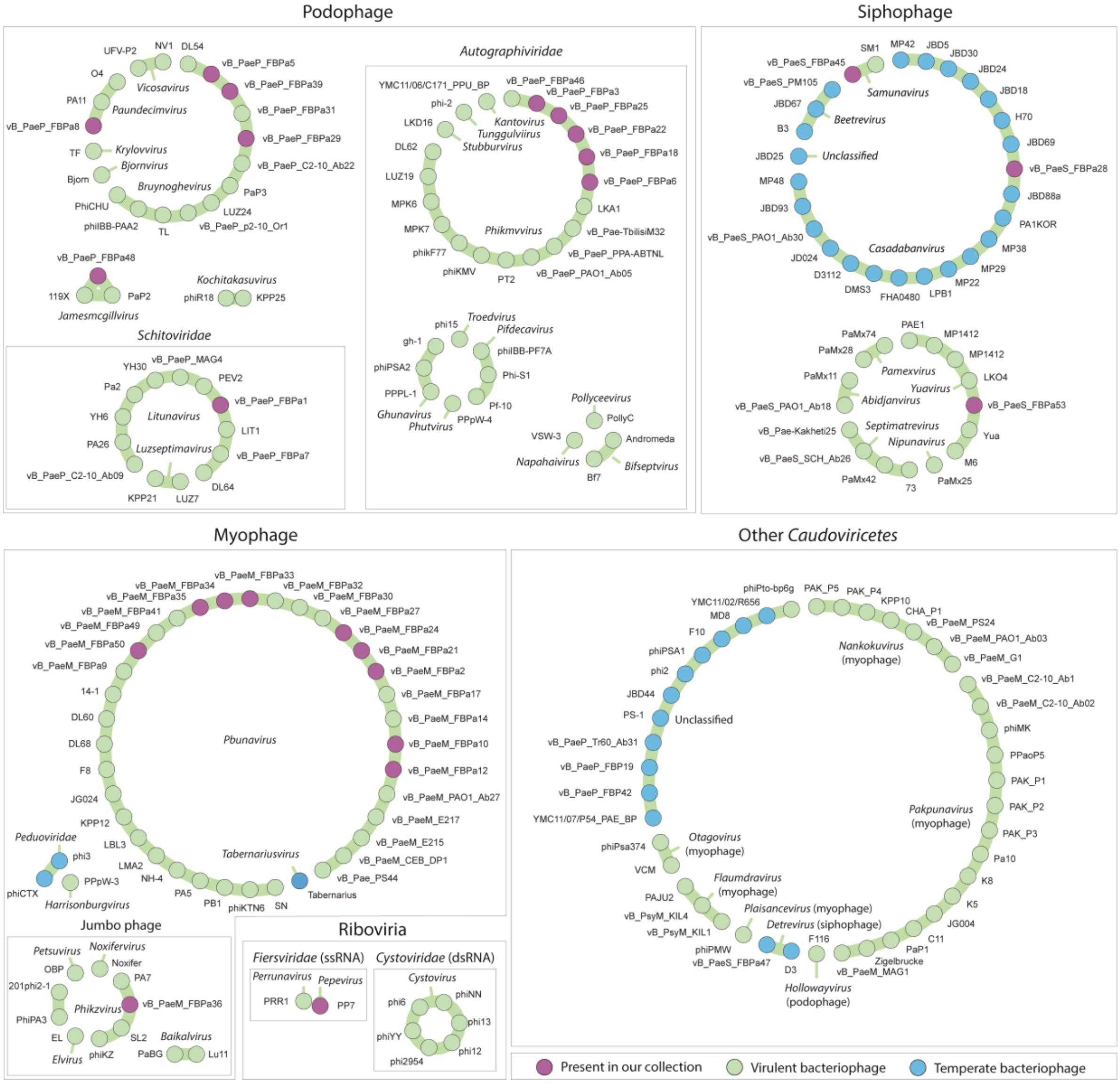


**Fig. S2.** Diversity of the phage panel used in this study. *Pseudomonas aeruginosa* phages were clustered using vConTACT2, based on guilt-by-contig-association classification of genomic sequences, and visualized in Cytoscape. The phage families and genus are indicated in italics, and the phages from the collection are highlighted in purple.

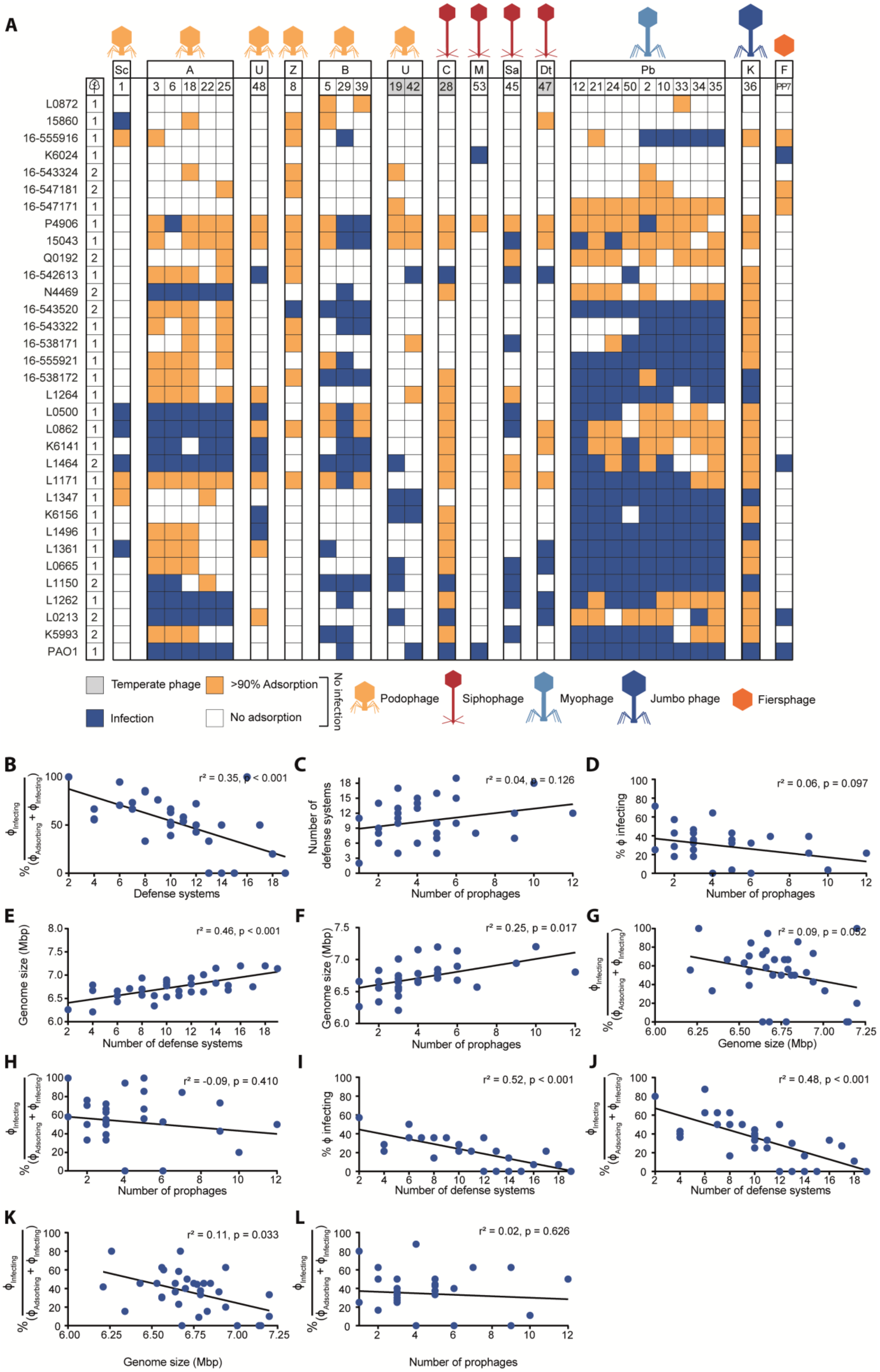
A
Sc
A
U
Z
B
U
C
M
Sa
Dt
Pb
K
F
1
3 6 18 22 25
48
8
5 29 39
19 42
28
53
45
47
12 21 24 50 2 10 33 34 35
36
PP7
L0872 1
15860 1
16-555916 1
K6024 1
16-543324 2
16-547181 2
16-547171 1
P4906 1
15043 1
Q0192 2
16-542613 1
N4469 2
16-543520 2
16-543322 1
16-538171 1
16-555921 1
16-538172 1
L1264 1
L0500 1
L0862 1
K6141 1
L1464 2
L1171 1
L1347 1
K6156 1
L1496 1
L1361 1
L0665 1
L1150 2
L1262 1
L0213 2
K5993 2
PAO1 1
Temperate phage
>90% Adsorption
Infection
No adsorption
No infection
Podophage
Siphophage
Myophage
Jumbo phage
Fiersphage
B
r² = 0.35, p < 0.001
Defense systems
C
r² = 0.04, p = 0.126
Number of defense systems
Number of prophages
D
r² = 0.06, p = 0.097
% ϕ infecting
Number of prophages
E
r² = 0.46, p < 0.001
Genome size (Mbp)
Number of defense systems
F
r² = 0.25, p = 0.017
Genome size (Mbp)
Number of prophages
G
r² = 0.09, p = 0.052
Genome size (Mbp)
H
r² = -0.09, p = 0.410
Number of prophages
I
r² = 0.52, p < 0.001
% ϕ infecting
Number of defense systems
J
r² = 0.48, p < 0.001
Number of defense systems
K
r² = 0.11, p = 0.033
Genome size (Mbp)
L
r² = 0.02, p = 0.626
Number of prophages

**Fig. S3. Linear regression analysis of correlation between different variables and phage resistance in *Pseudomonas aeruginosa*. (A)** Host range of phages against 32 clinical isolates of *P. aeruginosa* and strain PAO1. Phages are clustered by phylogeny. Phage-bacteria interactions are depicted as infection (blue), adsorption (>90%) but no infection (orange), or no interaction (white). Letters above the phage numbers indicate family or genus (for phages unassigned to a family): A, *Autographiviridae*; B, *Bruynoghevirus*; C, *Casadabanvirus*; Dt, *Detrevirus*; F, Fiersviridae; K, *Phikzvirus*; M, *Mesyanzhinovviridae*; Pb, *Pbunavirus*; Sa, *Samunavirus*; Sc, *Schitoviridae*; Z, *Zobellviridae*; U, unassigned. Linear regression analysis considering all 28 phages of the panel for **(B)** genome size and number of prophages; **(C)** number of defense systems and number of prophages; **(D)** percentage of infecting phages and number of prophages; **(E)** genome size and number of defense systems. Linear regression analysis considering all 28 phages and adsorption at a conservative level of 90% for **(F)** percentage of adsorbing phages that can establish a productive infection (% $\phi_{Infecting}$ / ($\phi_{Adsorbing}$+$\phi_{Infecting}$)) and number of defense systems; **(G)** percentage of adsorbing phages that can establish a productive infection and genome size; **(H)** percentage of absorbing phages that can establish a productive infection and number of prophages. Linear regression analysis considering a selection of representative phages for **(I)** percentage of infecting phages and number of defense systems; **(J)** percentage of adsorbing phages that can establish a productive infection (% $\phi_{Infecting}$ / ($\phi_{Adsorbing}$+$\phi_{Infecting}$)) and number of defense systems; **(K)** percentage of adsorbing phages that can establish a productive infection and genome size; **(L)** percentage of absorbing phages that can establish a productive infection and number of prophages. $r^2$ represents adjusted R-squared, a goodness-of-fit measure for the linear regression models. Representative phages include ϕPa1, ϕPa2, ϕPa8, ϕPa12, ϕPa28, ϕPa34, ϕPa36, ϕPa39, ϕPa42, ϕPa45, ϕPa47, ϕPa48, ϕPa53, and ϕPP7.

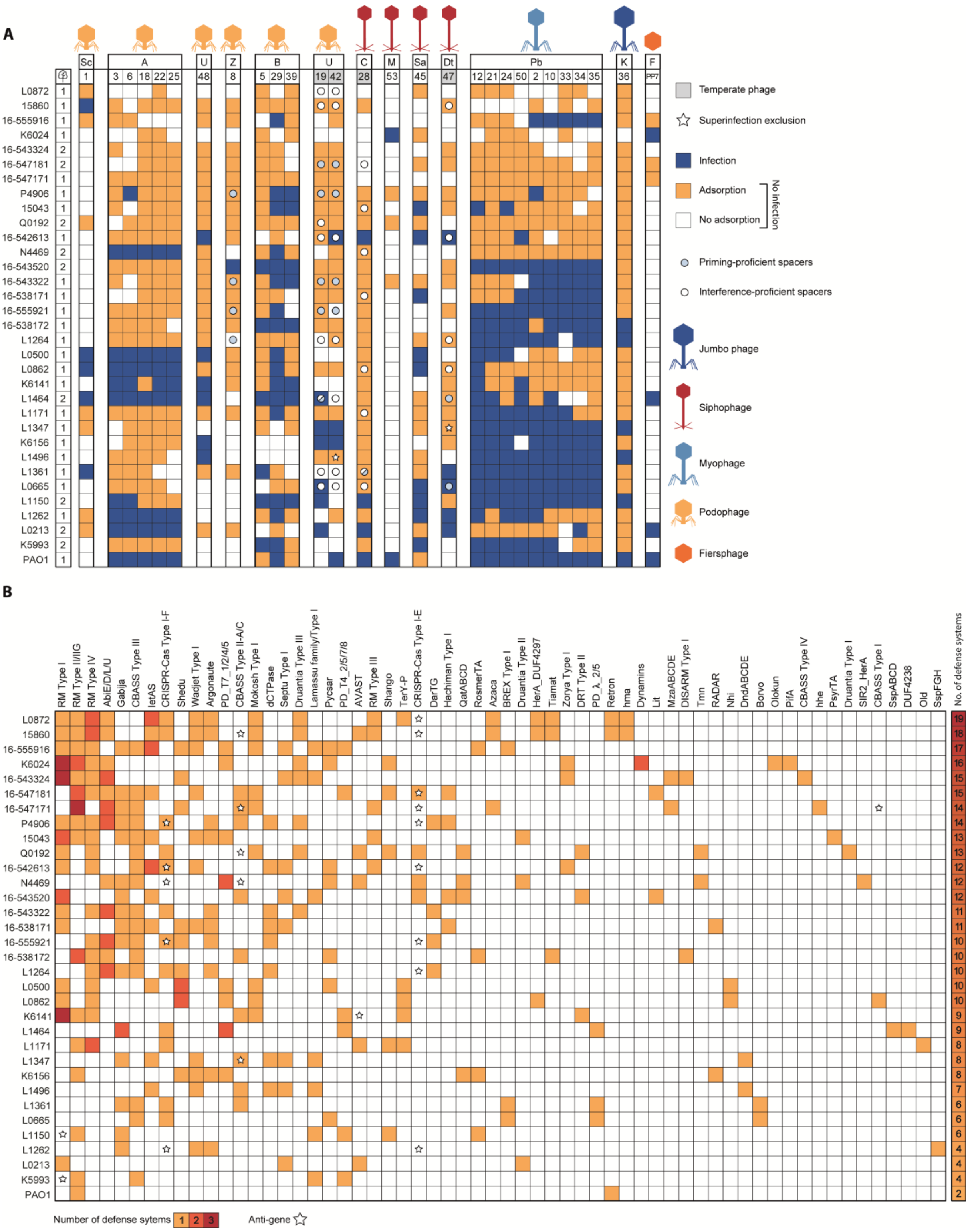


**Fig. S4. Role of adaptive immunity, superinfection exclusion, and anti-defense genes in defining the phage host range in *Pseudomonas aeruginosa*. (A)** Spacers found in the clinical isolates to match the phages in our panel. Spacers that have a matching protospacer adjacent motif (PAM) and protospacer are labelled as interference-proficient, indicated by white circles. Spacers with a ±1 slipped PAM or with up to 5 protospacer mutations are labelled as priming-proficient, indicated by blue circles. Superinfection exclusion occurrences are denoted with a star symbol when a strain carries a prophage that resembles a temperate phage in the phage panel, with a pident >90% and coverage >85%. Phage-bacteria interactions are depicted as infection (blue), adsorption but no infection (orange), or no interaction (white). Letters above the phage numbers indicate family or genus (for phages unassigned to a family): A, *Autographiviridae*; B, *Bruynoghevirus*; C, *Casadabanvirus*; Dt, *Detrevirus*; F, Fiersviridae; K, *Phikzvirus*; M, *Mesyanzhinovviridae*; Pb, *Pbunavirus*; Sa, *Samunavirus*; Sc, *Schitoviridae*; Z, *Zobellviridae*; U, unassigned. **(B)** Anti-defense genes found in the genomes of the clinical isolates are indicated with a star in the position corresponding to the defense system potentially affected by the anti-defense gene. The number of instances of each defense system type per strain is indicated in yellow, orange, or red for 1, 2, or 3 respectively. The total number of defense systems found per strain is indicated in a heatmap bar on the right. The complete list of spacers and anti-defense genes found in the clinical strains and phages can be found in **Table S4** and **Table S5**, respectively.

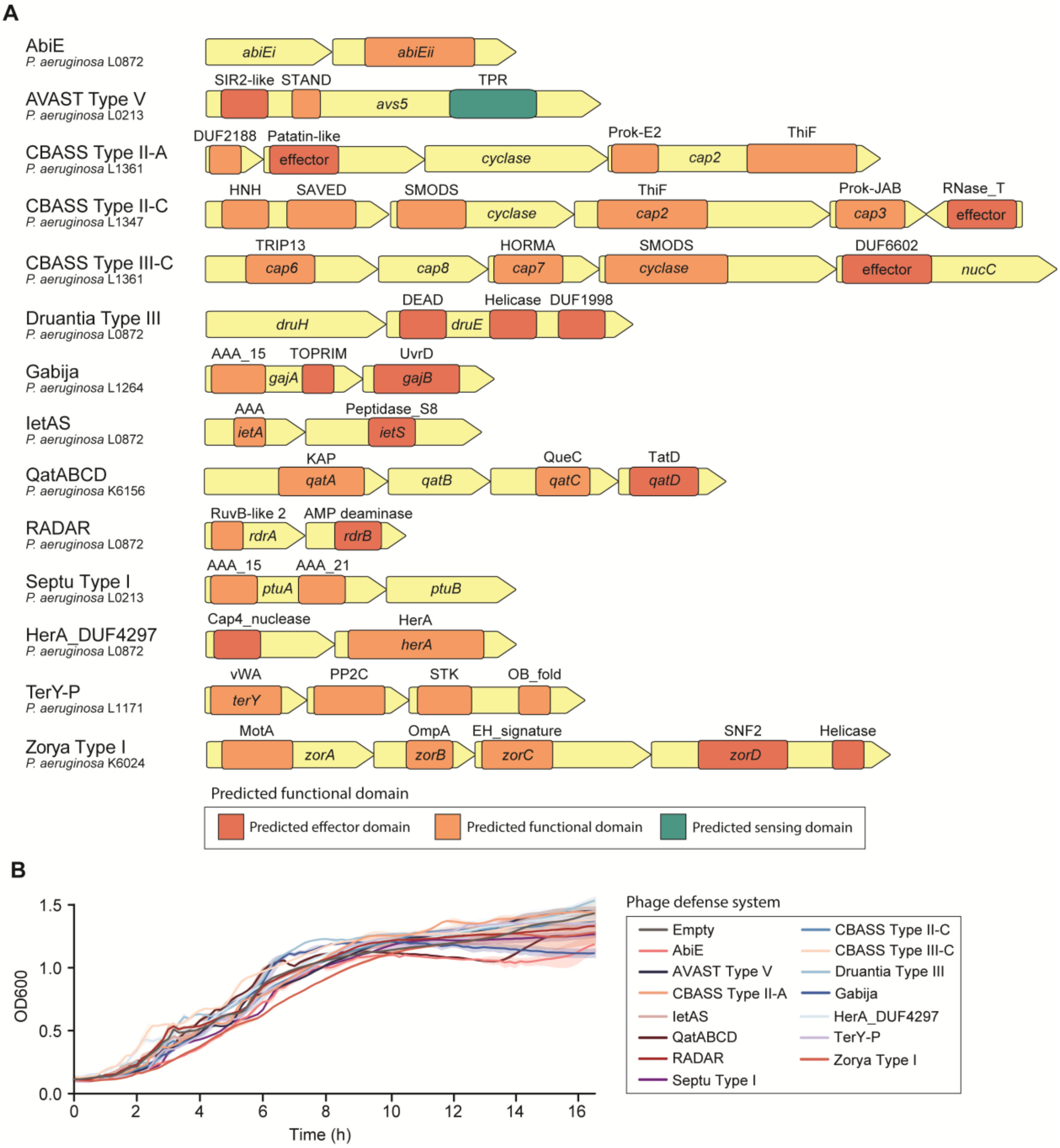


**Fig. S5. Individual defense systems cloned in *Pseudomonas aeruginosa* strain PAO1.** **(A)** Gene composition and functional domains of the 14 defense systems cloned into *P. aeruginosa* strain PAO1. **(B)** Growth of PAO1 strains containing individual defense systems. No toxic effect is observed in cell growth, as measured by optical density at 600 nm.

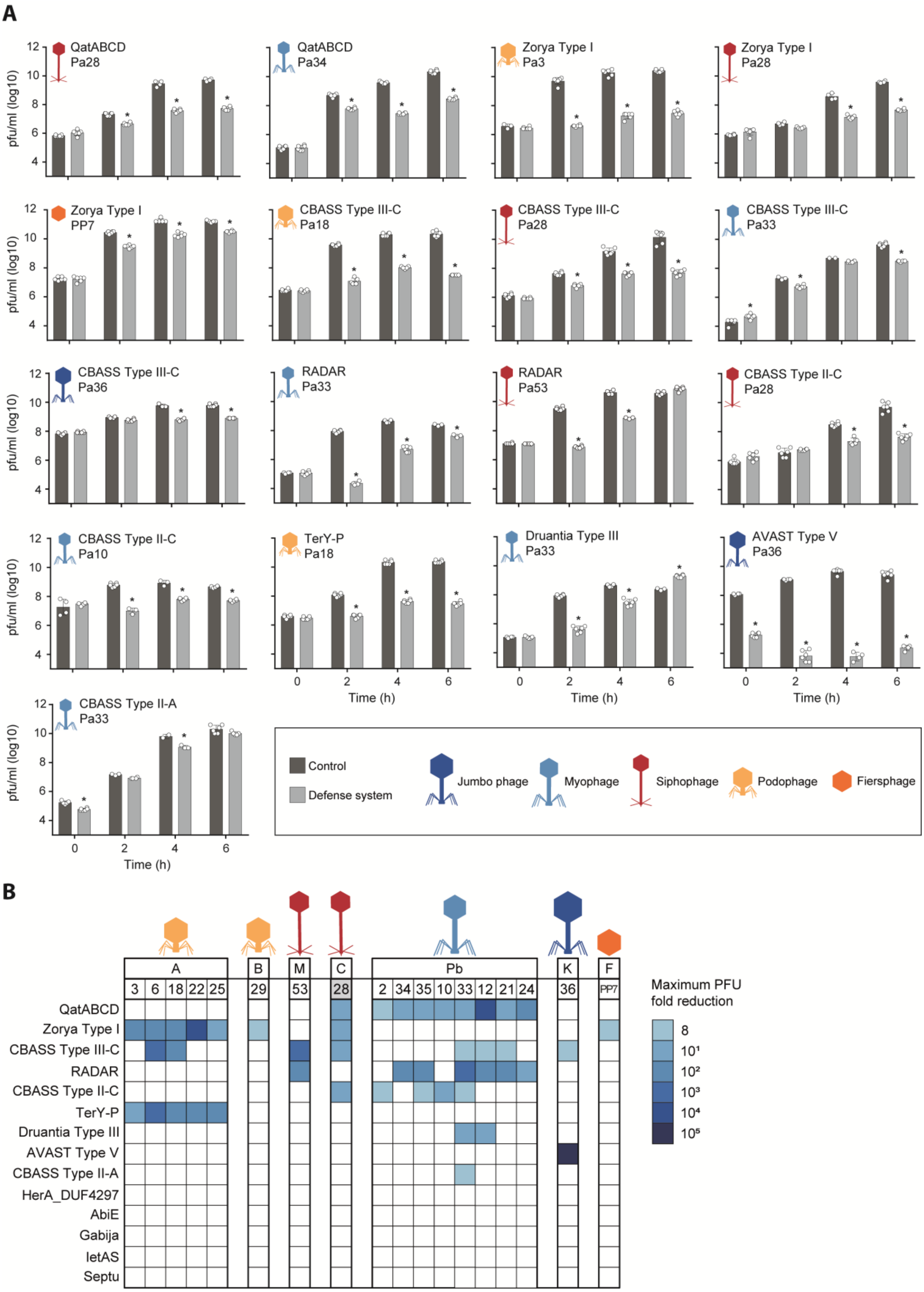


**Fig. S6. Effect of individual defense systems on phage infection dynamics in *Pseudomonas aeruginosa* PAO1. (A)** Graphical representation of the infection dynamics of phages representative of the families targeted by each defense system. Results are shown as the mean ± standard deviation of the phage concentration measured at 0, 2, 4, and 6 hours post-infection of the cells containing an empty plasmid (control) or defense system. Statistically significant differences ($p < 0.01$) were determined by two-way ANOVA followed by Sidak's multiple comparison test and are indicated with *. **(B)** Heatmap summarizing the results of the infection dynamics assay. The heatmap shows the maximum fold reduction in phage concentration obtained during the infection dynamics assay. Letters above the phage numbers indicate family or genus (for phages unassigned to a family): A, *Autographiviridae*; B, *Bruynoghevirus*; C, *Casadabanvirus*; F, Fiersviridae; K, *Phikzvirus*; M, *Mesyanzhinovviridae*; Pb, *Pbunavirus*; U, unassigned.

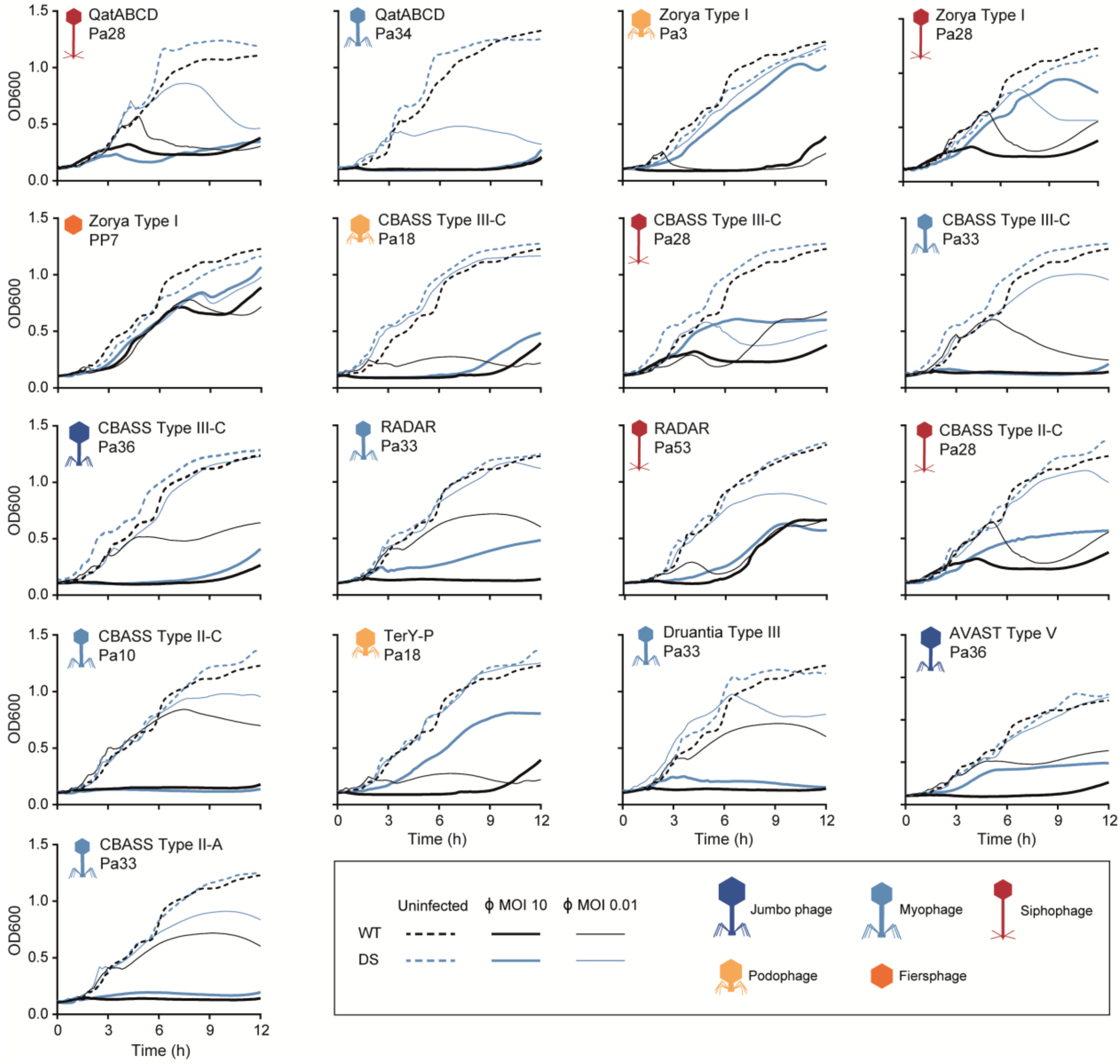


**Fig. S7. Effect of individual defense systems on bacterial growth during phage infection.** Graphical representation of the bacterial growth upon infection with phage representatives of the families targeted by each defense system. Results are shown as the average optical density at 600 nm of cultures of strains containing a control plasmid (WT) or defense system (DS), uninfected or infected with phage at a multiplicity of infection (MOI) of 10 or 0.01.

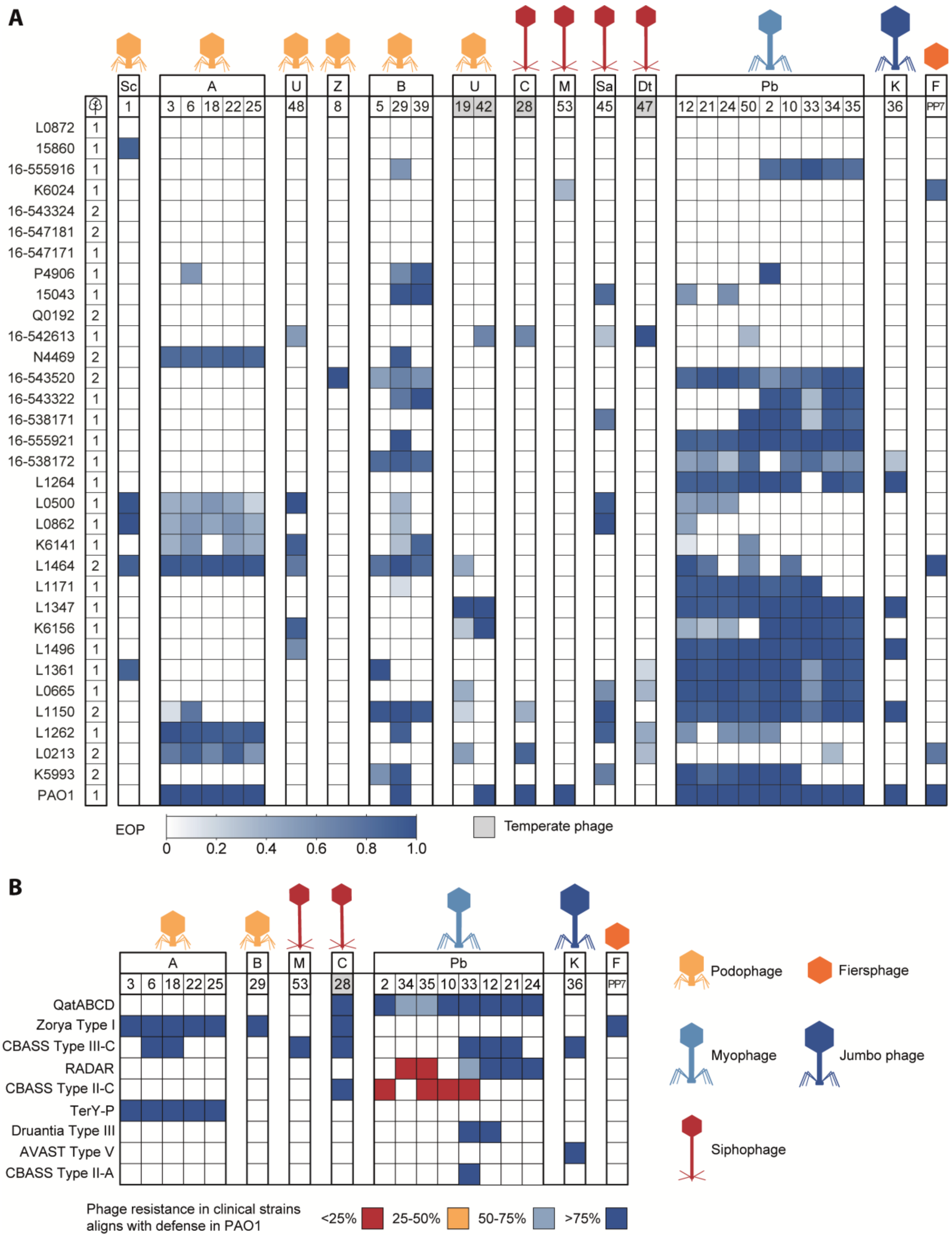


**Fig. S8. Predictive value of defense system activity in PAO1 to phage susceptibility of the clinical strains. (A)** Efficiency of platting (EOP) of 28 phages against the panel of 32 clinical strains and PAO1. The EOP was determined as the ratio of the phage concentration in each clinical strain to that in either PAO1 or the strain that the phage infects most efficiently. **(B)** Heatmap representation of matches between the phage infectivity profile in the clinical strains and the expected protection based on assays in PAO1 strains containing individual defense systems. The matches are categorized into four groups: <25%, 25-50%, 50-75%, and >75%, which are color-coded as red, yellow, light blue, and dark blue, respectively. Letters above the phage numbers indicate family or genus (for phages unassigned to a family): A, *Autographiviridae*; B, *Bruynoghevirus*; C, *Casadabanvirus*; Dt, *Detrevirus*; F, Fiersviridae; K, *Phikzvirus*; M, *Mesyanzhinovviridae*; Pb, *Pbunavirus*; Sa, *Samunavirus*; Sc, *Schitoviridae*; Z, *Zobellviridae*; U, unassigned.

**Table S1. (separate file)**

Matrix of defense systems identified in the 311 RefSeq genomes of *Pseudomonas aeruginosa*.

**Table S2. (separate file)**

Features of the clinical isolates of *Pseudomonas aeruginosa* used in this work, including defense system presence.

**Table S3. (separate file)**

Features of the *Pseudomonas aeruginosa* phages used in this study.

**Table S4. (separate file)**

List of CRISPR-Cas Type I-F and I-E interference-proficient and priming-proficient spacers found in the clinical isolates of *Pseudomonas aeruginosa* to target phages from our collection.

**Table S5. (separate file)**

Anti-defense genes found in our collection of *Pseudomonas aeruginosa* clinical strains and phages.

**Table S6. (separate file)**

Co-occurrence of defense systems in the *Pseudomonas aeruginosa* genomes of the RefSeq database identified by Coinfinder.

**Table S7. (separate file)**

List of primers used in this work.

**Table S8. (separate file)**

List of plasmids used in this work.
